# Ineffective neutralization of the SARS-CoV-2 Mu variant by convalescent and vaccine sera

**DOI:** 10.1101/2021.09.06.459005

**Authors:** Keiya Uriu, Izumi Kimura, Kotaro Shirakawa, Akifumi Takaori-Kondo, Taka-aki Nakada, Atsushi Kaneda, The Genotype to Phenotype Japan (G2P-Japan) Consortium, So Nakagawa, Kei Sato

## Abstract

On August 30, 2021, the WHO classified the SARS-CoV-2 Mu variant (B.1.621 lineage) as a new variant of interest. The WHO defines “comparative assessment of virus characteristics and public health risks” as primary action in response to the emergence of new SARS-CoV-2 variants. Here, we demonstrate that the Mu variant is highly resistant to sera from COVID-19 convalescents and BNT162b2-vaccinated individuals. Direct comparison of different SARS-CoV-2 spike proteins revealed that Mu spike is more resistant to serum-mediated neutralization than all other currently recognized variants of interest (VOI) and concern (VOC). This includes the Beta variant (B.1.351) that has been suggested to represent the most resistant variant to convalescent and vaccinated sera to date (e.g., Collier et al, Nature, 2021; Wang et al, Nature, 2021). Since breakthrough infection by newly emerging variants is a major concern during the current COVID-19 pandemic (Bergwerk et al., NEJM, 2021), we believe that our findings are of significant public health interest. Our results will help to better assess the risk posed by the Mu variant for vaccinated, previously infected and naïve populations.

## Text

During the current pandemic, severe acute respiratory syndrome coronavirus 2 (SARS-CoV-2), the causative agent of coronavirus disease 2019 (COVID-19), has considerably diversified. As of September 2021, the WHO has defined four variants of concern (VOC), Alpha (B.1.1.7), Beta (B.1.351), Gamma (P.1) and Delta (B.1.617.2 and AY lineages), as well as five variants of interest (VOI), Eta (B.1.525), Iota (B.1.526), Kappa (B.1.617.1), Lambda (C.37), and Mu (B.1.621).^1^

The Mu variant represents the most recently recognized VOI.^1^ Until August 30, 2021, this VOI was detected in 39 countries (**Table S1**). The epicenter of the Mu variant is Colombia, where it was first isolated on January 11, 2021 (GISAID ID: EPI_ISL_1220045; **Figure 1A** and **Table S2**). This country has experienced a huge COVID-19 surge from March to August 2021 that has peaked at 33,594 cases per day (on June 26, 2021; **Figure 1A**). Although the Gamma VOC was dominant during the initial phase, the Mu VOI outcompeted all other variants including the Gamma VOC in May 2021 and has driven the epidemic in Colombia since then (**Figure 1A**).

**Figure 1.**
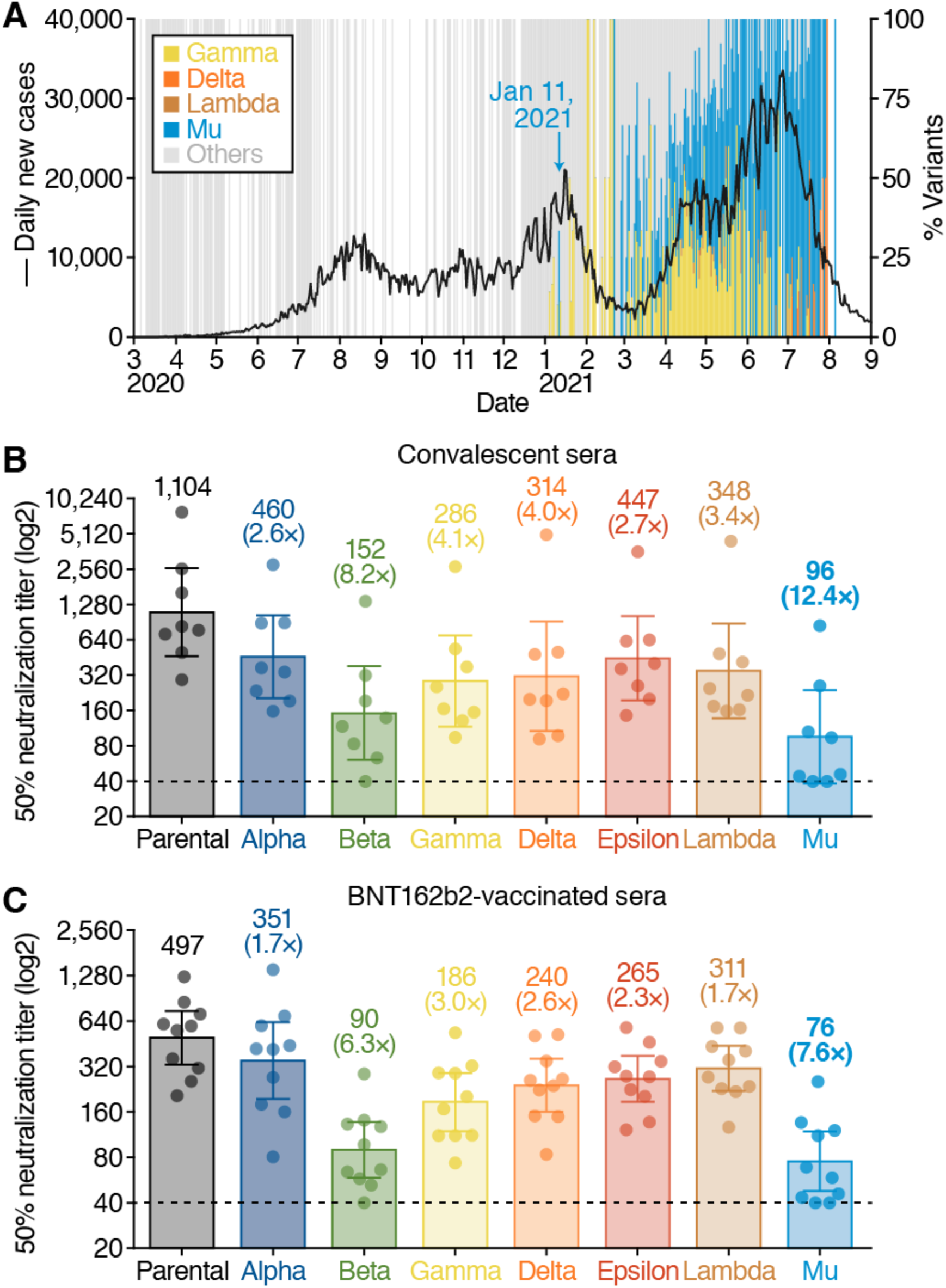
Characterization of the Mu variant. (**A**) SARS-CoV-2 epidemic in Colombia. New COVID-19 cases per day (black line, left y-axis) and percentage of different SARS-CoV-2 variants spreading in Colombia (right y-axis) are shown. The daily frequency of Gamma (P.1), Delta (B.1.617.2, AY.4, AY.5, AY.12), Lambda (C.37), Mu (B.1.621), and other variants are shown in the indicated colors. Note that there are a few Delta VOC (the currently most dominant variant in the world) and Lambda VOI (a variant mainly spreading in South American countries) have been isolated in this country so far. The date when the Mu variant was first isolated (January 11, 2021) is indicated in the figure. The raw data are summarized in **Table S2** in the Supplementary Appendix. (**B and C**) Virus neutralization assays. A neutralization assay was performed using pseudoviruses harboring the SARS-CoV-2 spike proteins of the Alpha, Beta, Gamma, Delta, Epsilon, Lambda, Mu variants or the D614G-harboring parental virus. Eight COVID-19 convalescent sera (**B**) and ten sera from BNT162b2-vaccinated individuals (**C**) were tested. The assay of each serum was performed in triplicate to determine the 50% neutralization titer, and each data point represents the 50% neutralization titer obtained with a serum sample against the indicated pseudovirus. The bar graphs indicate geometric mean titers with 95% confidence. The numbers over the bars indicate geometric mean titers. The numbers over the bars in parentheses (with “X”) indicate the average of fold change in neutralization resistance of the indicated spike variants compared to that with the parental spike in each serum. Statistical analysis was performed with the use of the Wilcoxon signed-rank test. Horizontal dashed lines indicate limit of detection. The raw data are summarized in **Tables S6 and S7** in the Supplementary Appendix.

Newly emerging SARS-CoV-2 variants need to be carefully monitored for a potential increase in transmission rate, pathogenicity and/or resistance to immune responses. For example, the resistance of VOC/VOIs to humoral immunity elicited by natural SARS-CoV-2 infection or vaccination may allow significant spread of the virus in populations that were initially thought to be protected.^2^ Resistance to COVID-19 convalescent and vaccine recipient sera can be attributed to a variety of mutations in the viral spike protein.^2^ The majority of Mu variants harbors the following eight mutations in spike: T95I, YY144-145TSN, R346K, E484K, N501Y, D614G, P681H, and D950N (**Tables S3 and S4**). These include mutations commonly identified in VOCs: E484K (shared with Beta, Gamma), N501Y (shared with Alpha), P681H (shared with Alpha) and D950N (shared with Delta) (**Table S5**). Of those, the E484K change has been shown to reduce sensitivity towards antibodies induced by natural SARS-CoV-2 infection and vaccination.^3,4^

To assess the sensitivity of the Mu variant to antibodies induced by SARS-CoV-2 infection and vaccination, we generated pseudoviruses harboring the spike proteins of Mu or the other VOC/VOIs. Virus neutralization assays revealed that the Mu variant is 12.4-fold more resistant to sera of eight COVID-19 convalescents, who were infected during the early pandemic (April–September, 2020), than the parental virus (*P*=0.0078; **Figure 1B**). Also, the Mu variant was 7.6-fold more resistant to sera obtained from ten BNT162b2-vaccinated individuals compared to the parental virus (*P*=0.0020; **Figure 1C**). Notably, although the Beta VOC was thought to be the most resistant variant to date,^3,4^ Mu pseudoviruses were significantly more resistant to convalescent serum-mediated neutralization than Beta pseudoviruses (*P*=0.031; **Figure 1B**). Thus, the Mu variant shows a pronounced resistance to antibodies elicited by natural SARS-CoV-2 infection and the BNT162b2 mRNA vaccine. Since breakthrough infections are a major threat of newly emerging SARS-CoV-2 variants,^5^ we strongly suggest to further characterize and monitor the Mu variant.

## Notes

**Conflict of interest:** The authors declare that no competing interests exist.

### Competing Interest Statement

The authors have declared no competing interest.

