## Supplementary Method and Tables for "Ineffective neutralization of the SARS-CoV-2 Mu variant by convalescent and vaccine sera"

### Supplementary Appendix

### Table of Contents


| | Page |
|---|---|
| **Materials and Methods** | 2 |
| Ethics statement | |
| Human sera | |
| Epidemic data | |
| Viral genome sequences | |
| Cell culture | |
| Plasmid construction | |
| Neutralization assay | |
| **Table S1.** Number of Mu variant sequences isolated in each country. | 4 |
| **Table S2.** Number and percentage of different VOCs/VOIs in Colombia. | 5-14 |
| **Table S3.** Haplotypes of the spike protein of Mu variant. | 15 |
| **Table S4.** Mutations in the spike protein of Mu variant. | 16 |
| **Table S5.** Mutations in the spike proteins of SARS-CoV-2 variants used in this study. | 17 |
| **Table S6.** Summary of COVID-19 convalescent sera used in this study. | 18 |
| **Table S7.** Summary of sera from BNT162b2-vaccinated individuals used in this study. | 19 |
| **Table S8.** Primers used for the construction of Mu spike expression plasmid. | 20 |
| **Consortia** | 21 |
| **Acknowledgments** | 22 |
| **Supplemental References** | 23 |

### Materials and Methods

#### Ethics statement

For the use of human specimens, all protocols involving human subjects recruited at Kyoto University and Chiba University were reviewed and approved by the Institutional Review Boards of Kyoto University (approval number G0697) and Chiba University (approval number HS202103-03). All human subjects provided written informed consent.

#### Human sera

Peripheral blood was collected from three COVID-19 convalescents and the sera were isolated (**Table S6**). Moreover, peripheral blood was collected four weeks after the second vaccination with BNT162b2 (Pfizer-BioNTech), and the sera of ten vaccinated-individuals were isolated (**Table S7**). Sera were inactivated at 56°C for 30 min and stored at –80°C until use. Five additional COVID-19 convalescent sera were purchased from RayBiotech (**Table S6**).

#### Epidemic data

The data of daily COVID-19 cases in Colombia (until August 31, 2021) were downloaded from Our World in Data (https://ourworldindata.org/covid-cases)[6] on September 2, 2021.

#### Viral genome sequences

SARS-CoV-2 annotation information used in this study was downloaded from the GISAID database (https://www.gisaid.org) as of August 30, 2021 (3,059,698 genomes). Based on the annotation by Phylogenetic Assignment of Named Global Outbreak (PANGO), we determined the daily frequency of SARS-CoV-2 Gamma (P.1), Delta (B.1.617.2, AY.4, AY.5, AY.12), Lambda (C.37), and Mu (B.1.621) variants isolated in Colombia. We also examined amino acid replacements of the spike protein of each SARS-CoV-2 variant by comparing it to the sequence of the Wuhan-Hu-1 strain (GISAID ID: EPI_ISL_1532199).

#### Cell culture

HEK293T cells (a human embryonic kidney cell line; ATCC CRL-3216) and HOS-ACE2/TMPRSS2 cells,[7,8] a derivative of HOS cells (a human osteosarcoma cell line; ATCC CRL-1543) stably expressing human ACE2 and TMPRSS2, were maintained in Dulbecco's modified Eagle's medium (high glucose) (Wako, Cat# 044-29765) containing 10% fetal calf serum, 100 units penicillin and 100 ug/ml streptomycin.

#### Plasmid construction

Plasmids expressing the SARS-CoV-2 spike proteins of the parental D614G (B.1), Alpha (B.1.1.7), Beta (B.1.351), Gamma (P.1), Delta (B.1.617.2), Epsilon (B.1.427), and Lambda (C.37) variants were prepared in our previous studies.[8-10] A plasmid expressing the spike protein of the Mu (B.1.621) variant was generated by site-directed overlap extension PCR using pC-SARS2-S D614G[8] as the template and the primers listed in **Table S8**. The resulting PCR fragment was digested with KpnI and NotI and inserted into the corresponding site of the pCAGGS vector.[11] Nucleotide sequences were determined by DNA sequencing services (Eurofins), and the sequence data were analyzed by Sequencher v5.1 software (Gene Codes Corporation).

**Neutralization assay**

Pseudoviruses were prepared as previously described.[8-10,12] Briefly, lentivirus (HIV-1)-based, luciferase-expressing reporter viruses were pseudotyped with the SARS-CoV-2 spikes. HEK293T cells ($1 \times 10^6$ cells) were cotransfected with 1 μg psPAX2-IN/HiBiT,[13] 1 μg pWPI-Luc2,[13] and 500 ng plasmids expressing parental S or its derivatives using PEI Max (Polysciences, Cat# 24765-1) according to the manufacturer's protocol. Two days post transfection, the culture supernatants were harvested and centrifuged. The pseudoviruses were stored at –80°C until use.

Neutralization assays were performed as previously described.[9,12] Briefly, the SARS-CoV-2 spike pseudoviruses (counting ~20,000 relative light units) were incubated with serially diluted (40-, 120-, 360-, 1,080-, 3,240-, 9,720-, and 29,160-fold dilution at the final concentration) heat-inactivated human sera at 37°C for 1 h. Pseudoviruses without sera were included as controls. Then, an 80 μl mixture of pseudovirus and serum was added to HOS-ACE2/TMPRSS2 cells (10,000 cells/50 μl) in a 96-well white plate. Two days post infection, the infected cells were lysed with a One-Glo luciferase assay system (Promega, Cat# E6130), and the luminescent signal was measured using a GloMax explorer multimode microplate reader 3500 (Promega). The assay of each serum was performed in triplicate, and the 50% neutralization titer was calculated using Prism 9 (GraphPad Software).

**Table S1. Number of Mu variant sequences isolated in each country.**

| Country | B.1.621 (Mu) | B.1.621.1 |
|---|---|---|
| USA | 1,492 | 505 |
| Colombia | 844 | 8 |
| Spain | 344 | 122 |
| Mexico | 338 | 6 |
| Ecuador | 168 | 0 |
| Aruba | 74 | 5 |
| Chile | 70 | 2 |
| Netherlands | 68 | 2 |
| Italy | 63 | 14 |
| United Kingdom | 48 | 35 |
| Costa Rica | 46 | 0 |
| Canada | 43 | 4 |
| Switzerland | 23 | 25 |
| Belgium | 22 | 12 |
| Portugal | 21 | 3 |
| Curacao | 14 | 5 |
| France | 14 | 1 |
| Brazil | 11 | 0 |
| Denmark | 7 | 0 |
| Germany | 7 | 7 |
| Peru | 6 | 0 |
| Poland | 6 | 0 |
| Bonaire | 5 | 1 |
| Venezuela | 5 | 0 |
| Finland | 3 | 0 |
| Sweden | 3 | 0 |
| Dominican Republic | 2 | 41 |
| Hong Kong | 2 | 0 |
| Ireland | 2 | 2 |
| Japan | 2 | 0 |
| Slovakia | 2 | 1 |
| Turkey | 2 | 0 |
| Austria | 1 | 48 |
| Romania | 1 | 0 |
| Sint Maarten | 1 | 1 |
| Liechtenstein | 1 | 0 |
| Turks and Caicos Islands | 1 | 0 |
| Luxembourg | 0 | 1 |
| Malta | 0 | 1 |

**Table S2. Number and percentage of different VOCs/VOIs in Colombia.**

| | Number | | | | | Percentage | | | | |
|---|---|---|---|---|---|---|---|---|---|---|
| Date | All | Gamma | Delta | Lambda | Mu | Gamma | Delta | Lambda | Mu | Others |
| 2020/3/6 | 1 | 0 | 0 | 0 | 0 | 0 | 0 | 0 | 0 | 100 |
| 2020/3/7 | 0 | 0 | 0 | 0 | 0 | 0 | 0 | 0 | 0 | 0 |
| 2020/3/8 | 0 | 0 | 0 | 0 | 0 | 0 | 0 | 0 | 0 | 0 |
| 2020/3/9 | 0 | 0 | 0 | 0 | 0 | 0 | 0 | 0 | 0 | 0 |
| 2020/3/10 | 2 | 0 | 0 | 0 | 0 | 0 | 0 | 0 | 0 | 100 |
| 2020/3/11 | 5 | 0 | 0 | 0 | 0 | 0 | 0 | 0 | 0 | 100 |
| 2020/3/12 | 3 | 0 | 0 | 0 | 0 | 0 | 0 | 0 | 0 | 100 |
| 2020/3/13 | 2 | 0 | 0 | 0 | 0 | 0 | 0 | 0 | 0 | 100 |
| 2020/3/14 | 7 | 0 | 0 | 0 | 0 | 0 | 0 | 0 | 0 | 100 |
| 2020/3/15 | 3 | 0 | 0 | 0 | 0 | 0 | 0 | 0 | 0 | 100 |
| 2020/3/16 | 5 | 0 | 0 | 0 | 0 | 0 | 0 | 0 | 0 | 100 |
| 2020/3/17 | 4 | 0 | 0 | 0 | 0 | 0 | 0 | 0 | 0 | 100 |
| 2020/3/18 | 5 | 0 | 0 | 0 | 0 | 0 | 0 | 0 | 0 | 100 |
| 2020/3/19 | 3 | 0 | 0 | 0 | 0 | 0 | 0 | 0 | 0 | 100 |
| 2020/3/20 | 3 | 0 | 0 | 0 | 0 | 0 | 0 | 0 | 0 | 100 |
| 2020/3/21 | 0 | 0 | 0 | 0 | 0 | 0 | 0 | 0 | 0 | 0 |
| 2020/3/22 | 2 | 0 | 0 | 0 | 0 | 0 | 0 | 0 | 0 | 100 |
| 2020/3/23 | 2 | 0 | 0 | 0 | 0 | 0 | 0 | 0 | 0 | 100 |
| 2020/3/24 | 1 | 0 | 0 | 0 | 0 | 0 | 0 | 0 | 0 | 100 |
| 2020/3/25 | 4 | 0 | 0 | 0 | 0 | 0 | 0 | 0 | 0 | 100 |
| 2020/3/26 | 6 | 0 | 0 | 0 | 0 | 0 | 0 | 0 | 0 | 100 |
| 2020/3/27 | 1 | 0 | 0 | 0 | 0 | 0 | 0 | 0 | 0 | 100 |
| 2020/3/28 | 9 | 0 | 0 | 0 | 0 | 0 | 0 | 0 | 0 | 100 |
| 2020/3/29 | 3 | 0 | 0 | 0 | 0 | 0 | 0 | 0 | 0 | 100 |
| 2020/3/30 | 8 | 0 | 0 | 0 | 0 | 0 | 0 | 0 | 0 | 100 |
| 2020/3/31 | 16 | 0 | 0 | 0 | 0 | 0 | 0 | 0 | 0 | 100 |
| 2020/4/1 | 16 | 0 | 0 | 0 | 0 | 0 | 0 | 0 | 0 | 100 |
| 2020/4/2 | 6 | 0 | 0 | 0 | 0 | 0 | 0 | 0 | 0 | 100 |
| 2020/4/3 | 9 | 0 | 0 | 0 | 0 | 0 | 0 | 0 | 0 | 100 |
| 2020/4/4 | 15 | 0 | 0 | 0 | 0 | 0 | 0 | 0 | 0 | 100 |
| 2020/4/5 | 5 | 0 | 0 | 0 | 0 | 0 | 0 | 0 | 0 | 100 |
| 2020/4/6 | 3 | 0 | 0 | 0 | 0 | 0 | 0 | 0 | 0 | 100 |
| 2020/4/7 | 2 | 0 | 0 | 0 | 0 | 0 | 0 | 0 | 0 | 100 |
| 2020/4/8 | 0 | 0 | 0 | 0 | 0 | 0 | 0 | 0 | 0 | 0 |
| 2020/4/9 | 3 | 0 | 0 | 0 | 0 | 0 | 0 | 0 | 0 | 100 |
| 2020/4/10 | 0 | 0 | 0 | 0 | 0 | 0 | 0 | 0 | 0 | 0 |
| 2020/4/11 | 0 | 0 | 0 | 0 | 0 | 0 | 0 | 0 | 0 | 0 |
| 2020/4/12 | 1 | 0 | 0 | 0 | 0 | 0 | 0 | 0 | 0 | 100 |
| 2020/4/13 | 0 | 0 | 0 | 0 | 0 | 0 | 0 | 0 | 0 | 0 |
| 2020/4/14 | 5 | 0 | 0 | 0 | 0 | 0 | 0 | 0 | 0 | 100 |
| 2020/4/15 | 2 | 0 | 0 | 0 | 0 | 0 | 0 | 0 | 0 | 100 |
| 2020/4/16 | 3 | 0 | 0 | 0 | 0 | 0 | 0 | 0 | 0 | 100 |
| 2020/4/17 | 4 | 0 | 0 | 0 | 0 | 0 | 0 | 0 | 0 | 100 |
| 2020/4/18 | 3 | 0 | 0 | 0 | 0 | 0 | 0 | 0 | 0 | 100 |
| 2020/4/19 | 0 | 0 | 0 | 0 | 0 | 0 | 0 | 0 | 0 | 0 |
| 2020/4/20 | 4 | 0 | 0 | 0 | 0 | 0 | 0 | 0 | 0 | 100 |
| 2020/4/21 | 0 | 0 | 0 | 0 | 0 | 0 | 0 | 0 | 0 | 0 |
| 2020/4/22 | 8 | 0 | 0 | 0 | 0 | 0 | 0 | 0 | 0 | 100 |
| 2020/4/23 | 2 | 0 | 0 | 0 | 0 | 0 | 0 | 0 | 0 | 100 |
| 2020/4/24 | 1 | 0 | 0 | 0 | 0 | 0 | 0 | 0 | 0 | 100 |
| 2020/4/25 | 1 | 0 | 0 | 0 | 0 | 0 | 0 | 0 | 0 | 100 |
| 2020/4/26 | 3 | 0 | 0 | 0 | 0 | 0 | 0 | 0 | 0 | 100 |
| 2020/4/27 | 1 | 0 | 0 | 0 | 0 | 0 | 0 | 0 | 0 | 100 |

| | | | | | | | | | | |
|---|---|---|---|---|---|---|---|---|---|---|
| 2020/4/28 | 5 | 0 | 0 | 0 | 0 | 0 | 0 | 0 | 0 | 100 |
| 2020/4/29 | 0 | 0 | 0 | 0 | 0 | 0 | 0 | 0 | 0 | 0 |
| 2020/4/30 | 10 | 0 | 0 | 0 | 0 | 0 | 0 | 0 | 0 | 100 |
| 2020/5/1 | 3 | 0 | 0 | 0 | 0 | 0 | 0 | 0 | 0 | 100 |
| 2020/5/2 | 8 | 0 | 0 | 0 | 0 | 0 | 0 | 0 | 0 | 100 |
| 2020/5/3 | 22 | 0 | 0 | 0 | 0 | 0 | 0 | 0 | 0 | 100 |
| 2020/5/4 | 28 | 0 | 0 | 0 | 0 | 0 | 0 | 0 | 0 | 100 |
| 2020/5/5 | 2 | 0 | 0 | 0 | 0 | 0 | 0 | 0 | 0 | 100 |
| 2020/5/6 | 1 | 0 | 0 | 0 | 0 | 0 | 0 | 0 | 0 | 100 |
| 2020/5/7 | 1 | 0 | 0 | 0 | 0 | 0 | 0 | 0 | 0 | 100 |
| 2020/5/8 | 0 | 0 | 0 | 0 | 0 | 0 | 0 | 0 | 0 | 0 |
| 2020/5/9 | 0 | 0 | 0 | 0 | 0 | 0 | 0 | 0 | 0 | 0 |
| 2020/5/10 | 0 | 0 | 0 | 0 | 0 | 0 | 0 | 0 | 0 | 0 |
| 2020/5/11 | 1 | 0 | 0 | 0 | 0 | 0 | 0 | 0 | 0 | 100 |
| 2020/5/12 | 0 | 0 | 0 | 0 | 0 | 0 | 0 | 0 | 0 | 0 |
| 2020/5/13 | 0 | 0 | 0 | 0 | 0 | 0 | 0 | 0 | 0 | 0 |
| 2020/5/14 | 0 | 0 | 0 | 0 | 0 | 0 | 0 | 0 | 0 | 0 |
| 2020/5/15 | 0 | 0 | 0 | 0 | 0 | 0 | 0 | 0 | 0 | 0 |
| 2020/5/16 | 0 | 0 | 0 | 0 | 0 | 0 | 0 | 0 | 0 | 0 |
| 2020/5/17 | 0 | 0 | 0 | 0 | 0 | 0 | 0 | 0 | 0 | 0 |
| 2020/5/18 | 0 | 0 | 0 | 0 | 0 | 0 | 0 | 0 | 0 | 0 |
| 2020/5/19 | 1 | 0 | 0 | 0 | 0 | 0 | 0 | 0 | 0 | 100 |
| 2020/5/20 | 0 | 0 | 0 | 0 | 0 | 0 | 0 | 0 | 0 | 0 |
| 2020/5/21 | 0 | 0 | 0 | 0 | 0 | 0 | 0 | 0 | 0 | 0 |
| 2020/5/22 | 0 | 0 | 0 | 0 | 0 | 0 | 0 | 0 | 0 | 0 |
| 2020/5/23 | 0 | 0 | 0 | 0 | 0 | 0 | 0 | 0 | 0 | 0 |
| 2020/5/24 | 0 | 0 | 0 | 0 | 0 | 0 | 0 | 0 | 0 | 0 |
| 2020/5/25 | 1 | 0 | 0 | 0 | 0 | 0 | 0 | 0 | 0 | 100 |
| 2020/5/26 | 0 | 0 | 0 | 0 | 0 | 0 | 0 | 0 | 0 | 0 |
| 2020/5/27 | 0 | 0 | 0 | 0 | 0 | 0 | 0 | 0 | 0 | 0 |
| 2020/5/28 | 1 | 0 | 0 | 0 | 0 | 0 | 0 | 0 | 0 | 100 |
| 2020/5/29 | 1 | 0 | 0 | 0 | 0 | 0 | 0 | 0 | 0 | 100 |
| 2020/5/30 | 0 | 0 | 0 | 0 | 0 | 0 | 0 | 0 | 0 | 0 |
| 2020/5/31 | 0 | 0 | 0 | 0 | 0 | 0 | 0 | 0 | 0 | 0 |
| 2020/6/1 | 0 | 0 | 0 | 0 | 0 | 0 | 0 | 0 | 0 | 0 |
| 2020/6/2 | 1 | 0 | 0 | 0 | 0 | 0 | 0 | 0 | 0 | 100 |
| 2020/6/3 | 1 | 0 | 0 | 0 | 0 | 0 | 0 | 0 | 0 | 100 |
| 2020/6/4 | 2 | 0 | 0 | 0 | 0 | 0 | 0 | 0 | 0 | 100 |
| 2020/6/5 | 0 | 0 | 0 | 0 | 0 | 0 | 0 | 0 | 0 | 0 |
| 2020/6/6 | 1 | 0 | 0 | 0 | 0 | 0 | 0 | 0 | 0 | 100 |
| 2020/6/7 | 0 | 0 | 0 | 0 | 0 | 0 | 0 | 0 | 0 | 0 |
| 2020/6/8 | 1 | 0 | 0 | 0 | 0 | 0 | 0 | 0 | 0 | 100 |
| 2020/6/9 | 0 | 0 | 0 | 0 | 0 | 0 | 0 | 0 | 0 | 0 |
| 2020/6/10 | 1 | 0 | 0 | 0 | 0 | 0 | 0 | 0 | 0 | 100 |
| 2020/6/11 | 0 | 0 | 0 | 0 | 0 | 0 | 0 | 0 | 0 | 0 |
| 2020/6/12 | 0 | 0 | 0 | 0 | 0 | 0 | 0 | 0 | 0 | 0 |
| 2020/6/13 | 0 | 0 | 0 | 0 | 0 | 0 | 0 | 0 | 0 | 0 |
| 2020/6/14 | 0 | 0 | 0 | 0 | 0 | 0 | 0 | 0 | 0 | 0 |
| 2020/6/15 | 1 | 0 | 0 | 0 | 0 | 0 | 0 | 0 | 0 | 100 |
| 2020/6/16 | 5 | 0 | 0 | 0 | 0 | 0 | 0 | 0 | 0 | 100 |
| 2020/6/17 | 3 | 0 | 0 | 0 | 0 | 0 | 0 | 0 | 0 | 100 |
| 2020/6/18 | 1 | 0 | 0 | 0 | 0 | 0 | 0 | 0 | 0 | 100 |
| 2020/6/19 | 0 | 0 | 0 | 0 | 0 | 0 | 0 | 0 | 0 | 0 |
| 2020/6/20 | 1 | 0 | 0 | 0 | 0 | 0 | 0 | 0 | 0 | 100 |
| 2020/6/21 | 0 | 0 | 0 | 0 | 0 | 0 | 0 | 0 | 0 | 0 |
| 2020/6/22 | 4 | 0 | 0 | 0 | 0 | 0 | 0 | 0 | 0 | 100 |
| 2020/6/23 | 4 | 0 | 0 | 0 | 0 | 0 | 0 | 0 | 0 | 100 |

| | | | | | | | | | | |
|---|---|---|---|---|---|---|---|---|---|---|
| 2020/6/24 | 3 | 0 | 0 | 0 | 0 | 0 | 0 | 0 | 0 | 100 |
| 2020/6/25 | 1 | 0 | 0 | 0 | 0 | 0 | 0 | 0 | 0 | 100 |
| 2020/6/26 | 2 | 0 | 0 | 0 | 0 | 0 | 0 | 0 | 0 | 100 |
| 2020/6/27 | 2 | 0 | 0 | 0 | 0 | 0 | 0 | 0 | 0 | 100 |
| 2020/6/28 | 0 | 0 | 0 | 0 | 0 | 0 | 0 | 0 | 0 | 0 |
| 2020/6/29 | 0 | 0 | 0 | 0 | 0 | 0 | 0 | 0 | 0 | 0 |
| 2020/6/30 | 2 | 0 | 0 | 0 | 0 | 0 | 0 | 0 | 0 | 100 |
| 2020/7/1 | 1 | 0 | 0 | 0 | 0 | 0 | 0 | 0 | 0 | 100 |
| 2020/7/2 | 2 | 0 | 0 | 0 | 0 | 0 | 0 | 0 | 0 | 100 |
| 2020/7/3 | 1 | 0 | 0 | 0 | 0 | 0 | 0 | 0 | 0 | 100 |
| 2020/7/4 | 1 | 0 | 0 | 0 | 0 | 0 | 0 | 0 | 0 | 100 |
| 2020/7/5 | 3 | 0 | 0 | 0 | 0 | 0 | 0 | 0 | 0 | 100 |
| 2020/7/6 | 1 | 0 | 0 | 0 | 0 | 0 | 0 | 0 | 0 | 100 |
| 2020/7/7 | 7 | 0 | 0 | 0 | 0 | 0 | 0 | 0 | 0 | 100 |
| 2020/7/8 | 1 | 0 | 0 | 0 | 0 | 0 | 0 | 0 | 0 | 100 |
| 2020/7/9 | 2 | 0 | 0 | 0 | 0 | 0 | 0 | 0 | 0 | 100 |
| 2020/7/10 | 1 | 0 | 0 | 0 | 0 | 0 | 0 | 0 | 0 | 100 |
| 2020/7/11 | 1 | 0 | 0 | 0 | 0 | 0 | 0 | 0 | 0 | 100 |
| 2020/7/12 | 0 | 0 | 0 | 0 | 0 | 0 | 0 | 0 | 0 | 0 |
| 2020/7/13 | 5 | 0 | 0 | 0 | 0 | 0 | 0 | 0 | 0 | 100 |
| 2020/7/14 | 0 | 0 | 0 | 0 | 0 | 0 | 0 | 0 | 0 | 0 |
| 2020/7/15 | 4 | 0 | 0 | 0 | 0 | 0 | 0 | 0 | 0 | 100 |
| 2020/7/16 | 0 | 0 | 0 | 0 | 0 | 0 | 0 | 0 | 0 | 0 |
| 2020/7/17 | 2 | 0 | 0 | 0 | 0 | 0 | 0 | 0 | 0 | 100 |
| 2020/7/18 | 0 | 0 | 0 | 0 | 0 | 0 | 0 | 0 | 0 | 0 |
| 2020/7/19 | 0 | 0 | 0 | 0 | 0 | 0 | 0 | 0 | 0 | 0 |
| 2020/7/20 | 1 | 0 | 0 | 0 | 0 | 0 | 0 | 0 | 0 | 100 |
| 2020/7/21 | 0 | 0 | 0 | 0 | 0 | 0 | 0 | 0 | 0 | 0 |
| 2020/7/22 | 0 | 0 | 0 | 0 | 0 | 0 | 0 | 0 | 0 | 0 |
| 2020/7/23 | 1 | 0 | 0 | 0 | 0 | 0 | 0 | 0 | 0 | 100 |
| 2020/7/24 | 0 | 0 | 0 | 0 | 0 | 0 | 0 | 0 | 0 | 0 |
| 2020/7/25 | 1 | 0 | 0 | 0 | 0 | 0 | 0 | 0 | 0 | 100 |
| 2020/7/26 | 1 | 0 | 0 | 0 | 0 | 0 | 0 | 0 | 0 | 100 |
| 2020/7/27 | 1 | 0 | 0 | 0 | 0 | 0 | 0 | 0 | 0 | 100 |
| 2020/7/28 | 0 | 0 | 0 | 0 | 0 | 0 | 0 | 0 | 0 | 0 |
| 2020/7/29 | 0 | 0 | 0 | 0 | 0 | 0 | 0 | 0 | 0 | 0 |
| 2020/7/30 | 1 | 0 | 0 | 0 | 0 | 0 | 0 | 0 | 0 | 100 |
| 2020/7/31 | 0 | 0 | 0 | 0 | 0 | 0 | 0 | 0 | 0 | 0 |
| 2020/8/1 | 0 | 0 | 0 | 0 | 0 | 0 | 0 | 0 | 0 | 0 |
| 2020/8/2 | 0 | 0 | 0 | 0 | 0 | 0 | 0 | 0 | 0 | 0 |
| 2020/8/3 | 0 | 0 | 0 | 0 | 0 | 0 | 0 | 0 | 0 | 0 |
| 2020/8/4 | 1 | 0 | 0 | 0 | 0 | 0 | 0 | 0 | 0 | 100 |
| 2020/8/5 | 0 | 0 | 0 | 0 | 0 | 0 | 0 | 0 | 0 | 0 |
| 2020/8/6 | 1 | 0 | 0 | 0 | 0 | 0 | 0 | 0 | 0 | 100 |
| 2020/8/7 | 2 | 0 | 0 | 0 | 0 | 0 | 0 | 0 | 0 | 100 |
| 2020/8/8 | 7 | 0 | 0 | 0 | 0 | 0 | 0 | 0 | 0 | 100 |
| 2020/8/9 | 0 | 0 | 0 | 0 | 0 | 0 | 0 | 0 | 0 | 0 |
| 2020/8/10 | 0 | 0 | 0 | 0 | 0 | 0 | 0 | 0 | 0 | 0 |
| 2020/8/11 | 0 | 0 | 0 | 0 | 0 | 0 | 0 | 0 | 0 | 0 |
| 2020/8/12 | 1 | 0 | 0 | 0 | 0 | 0 | 0 | 0 | 0 | 100 |
| 2020/8/13 | 1 | 0 | 0 | 0 | 0 | 0 | 0 | 0 | 0 | 100 |
| 2020/8/14 | 0 | 0 | 0 | 0 | 0 | 0 | 0 | 0 | 0 | 0 |
| 2020/8/15 | 0 | 0 | 0 | 0 | 0 | 0 | 0 | 0 | 0 | 0 |
| 2020/8/16 | 0 | 0 | 0 | 0 | 0 | 0 | 0 | 0 | 0 | 0 |
| 2020/8/17 | 0 | 0 | 0 | 0 | 0 | 0 | 0 | 0 | 0 | 0 |
| 2020/8/18 | 3 | 0 | 0 | 0 | 0 | 0 | 0 | 0 | 0 | 100 |
| 2020/8/19 | 0 | 0 | 0 | 0 | 0 | 0 | 0 | 0 | 0 | 0 |

| | | | | | | | | | | |
|---|---|---|---|---|---|---|---|---|---|---|
| 2020/8/20 | 1 | 0 | 0 | 0 | 0 | 0 | 0 | 0 | 0 | 100 |
| 2020/8/21 | 0 | 0 | 0 | 0 | 0 | 0 | 0 | 0 | 0 | 0 |
| 2020/8/22 | 0 | 0 | 0 | 0 | 0 | 0 | 0 | 0 | 0 | 0 |
| 2020/8/23 | 0 | 0 | 0 | 0 | 0 | 0 | 0 | 0 | 0 | 0 |
| 2020/8/24 | 0 | 0 | 0 | 0 | 0 | 0 | 0 | 0 | 0 | 0 |
| 2020/8/25 | 0 | 0 | 0 | 0 | 0 | 0 | 0 | 0 | 0 | 0 |
| 2020/8/26 | 1 | 0 | 0 | 0 | 0 | 0 | 0 | 0 | 0 | 100 |
| 2020/8/27 | 0 | 0 | 0 | 0 | 0 | 0 | 0 | 0 | 0 | 0 |
| 2020/8/28 | 0 | 0 | 0 | 0 | 0 | 0 | 0 | 0 | 0 | 0 |
| 2020/8/29 | 0 | 0 | 0 | 0 | 0 | 0 | 0 | 0 | 0 | 0 |
| 2020/8/30 | 0 | 0 | 0 | 0 | 0 | 0 | 0 | 0 | 0 | 0 |
| 2020/8/31 | 0 | 0 | 0 | 0 | 0 | 0 | 0 | 0 | 0 | 0 |
| 2020/9/1 | 0 | 0 | 0 | 0 | 0 | 0 | 0 | 0 | 0 | 0 |
| 2020/9/2 | 1 | 0 | 0 | 0 | 0 | 0 | 0 | 0 | 0 | 100 |
| 2020/9/3 | 0 | 0 | 0 | 0 | 0 | 0 | 0 | 0 | 0 | 0 |
| 2020/9/4 | 1 | 0 | 0 | 0 | 0 | 0 | 0 | 0 | 0 | 100 |
| 2020/9/5 | 0 | 0 | 0 | 0 | 0 | 0 | 0 | 0 | 0 | 0 |
| 2020/9/6 | 0 | 0 | 0 | 0 | 0 | 0 | 0 | 0 | 0 | 0 |
| 2020/9/7 | 0 | 0 | 0 | 0 | 0 | 0 | 0 | 0 | 0 | 0 |
| 2020/9/8 | 0 | 0 | 0 | 0 | 0 | 0 | 0 | 0 | 0 | 0 |
| 2020/9/9 | 11 | 0 | 0 | 0 | 0 | 0 | 0 | 0 | 0 | 100 |
| 2020/9/10 | 0 | 0 | 0 | 0 | 0 | 0 | 0 | 0 | 0 | 0 |
| 2020/9/11 | 0 | 0 | 0 | 0 | 0 | 0 | 0 | 0 | 0 | 0 |
| 2020/9/12 | 0 | 0 | 0 | 0 | 0 | 0 | 0 | 0 | 0 | 0 |
| 2020/9/13 | 0 | 0 | 0 | 0 | 0 | 0 | 0 | 0 | 0 | 0 |
| 2020/9/14 | 0 | 0 | 0 | 0 | 0 | 0 | 0 | 0 | 0 | 0 |
| 2020/9/15 | 0 | 0 | 0 | 0 | 0 | 0 | 0 | 0 | 0 | 0 |
| 2020/9/16 | 0 | 0 | 0 | 0 | 0 | 0 | 0 | 0 | 0 | 0 |
| 2020/9/17 | 0 | 0 | 0 | 0 | 0 | 0 | 0 | 0 | 0 | 0 |
| 2020/9/18 | 0 | 0 | 0 | 0 | 0 | 0 | 0 | 0 | 0 | 0 |
| 2020/9/19 | 0 | 0 | 0 | 0 | 0 | 0 | 0 | 0 | 0 | 0 |
| 2020/9/20 | 0 | 0 | 0 | 0 | 0 | 0 | 0 | 0 | 0 | 0 |
| 2020/9/21 | 0 | 0 | 0 | 0 | 0 | 0 | 0 | 0 | 0 | 0 |
| 2020/9/22 | 3 | 0 | 0 | 0 | 0 | 0 | 0 | 0 | 0 | 100 |
| 2020/9/23 | 0 | 0 | 0 | 0 | 0 | 0 | 0 | 0 | 0 | 0 |
| 2020/9/24 | 0 | 0 | 0 | 0 | 0 | 0 | 0 | 0 | 0 | 0 |
| 2020/9/25 | 0 | 0 | 0 | 0 | 0 | 0 | 0 | 0 | 0 | 0 |
| 2020/9/26 | 0 | 0 | 0 | 0 | 0 | 0 | 0 | 0 | 0 | 0 |
| 2020/9/27 | 0 | 0 | 0 | 0 | 0 | 0 | 0 | 0 | 0 | 0 |
| 2020/9/28 | 0 | 0 | 0 | 0 | 0 | 0 | 0 | 0 | 0 | 0 |
| 2020/9/29 | 0 | 0 | 0 | 0 | 0 | 0 | 0 | 0 | 0 | 0 |
| 2020/9/30 | 0 | 0 | 0 | 0 | 0 | 0 | 0 | 0 | 0 | 0 |
| 2020/10/1 | 0 | 0 | 0 | 0 | 0 | 0 | 0 | 0 | 0 | 0 |
| 2020/10/2 | 0 | 0 | 0 | 0 | 0 | 0 | 0 | 0 | 0 | 0 |
| 2020/10/3 | 0 | 0 | 0 | 0 | 0 | 0 | 0 | 0 | 0 | 0 |
| 2020/10/4 | 1 | 0 | 0 | 0 | 0 | 0 | 0 | 0 | 0 | 100 |
| 2020/10/5 | 3 | 0 | 0 | 0 | 0 | 0 | 0 | 0 | 0 | 100 |
| 2020/10/6 | 0 | 0 | 0 | 0 | 0 | 0 | 0 | 0 | 0 | 0 |
| 2020/10/7 | 0 | 0 | 0 | 0 | 0 | 0 | 0 | 0 | 0 | 0 |
| 2020/10/8 | 0 | 0 | 0 | 0 | 0 | 0 | 0 | 0 | 0 | 0 |
| 2020/10/9 | 0 | 0 | 0 | 0 | 0 | 0 | 0 | 0 | 0 | 0 |
| 2020/10/10 | 14 | 0 | 0 | 0 | 0 | 0 | 0 | 0 | 0 | 100 |
| 2020/10/11 | 1 | 0 | 0 | 0 | 0 | 0 | 0 | 0 | 0 | 100 |
| 2020/10/12 | 0 | 0 | 0 | 0 | 0 | 0 | 0 | 0 | 0 | 0 |
| 2020/10/13 | 0 | 0 | 0 | 0 | 0 | 0 | 0 | 0 | 0 | 0 |
| 2020/10/14 | 0 | 0 | 0 | 0 | 0 | 0 | 0 | 0 | 0 | 0 |
| 2020/10/15 | 0 | 0 | 0 | 0 | 0 | 0 | 0 | 0 | 0 | 0 |

| | | | | | | | | | | |
|---|---|---|---|---|---|---|---|---|---|---|
| 2020/10/16 | 0 | 0 | 0 | 0 | 0 | 0 | 0 | 0 | 0 | 0 |
| 2020/10/17 | 1 | 0 | 0 | 0 | 0 | 0 | 0 | 0 | 0 | 100 |
| 2020/10/18 | 0 | 0 | 0 | 0 | 0 | 0 | 0 | 0 | 0 | 0 |
| 2020/10/19 | 0 | 0 | 0 | 0 | 0 | 0 | 0 | 0 | 0 | 0 |
| 2020/10/20 | 3 | 0 | 0 | 0 | 0 | 0 | 0 | 0 | 0 | 100 |
| 2020/10/21 | 1 | 0 | 0 | 0 | 0 | 0 | 0 | 0 | 0 | 100 |
| 2020/10/22 | 1 | 0 | 0 | 0 | 0 | 0 | 0 | 0 | 0 | 100 |
| 2020/10/23 | 0 | 0 | 0 | 0 | 0 | 0 | 0 | 0 | 0 | 0 |
| 2020/10/24 | 0 | 0 | 0 | 0 | 0 | 0 | 0 | 0 | 0 | 0 |
| 2020/10/25 | 1 | 0 | 0 | 0 | 0 | 0 | 0 | 0 | 0 | 100 |
| 2020/10/26 | 0 | 0 | 0 | 0 | 0 | 0 | 0 | 0 | 0 | 0 |
| 2020/10/27 | 1 | 0 | 0 | 0 | 0 | 0 | 0 | 0 | 0 | 100 |
| 2020/10/28 | 0 | 0 | 0 | 0 | 0 | 0 | 0 | 0 | 0 | 0 |
| 2020/10/29 | 0 | 0 | 0 | 0 | 0 | 0 | 0 | 0 | 0 | 0 |
| 2020/10/30 | 0 | 0 | 0 | 0 | 0 | 0 | 0 | 0 | 0 | 0 |
| 2020/10/31 | 2 | 0 | 0 | 0 | 0 | 0 | 0 | 0 | 0 | 100 |
| 2020/11/1 | 0 | 0 | 0 | 0 | 0 | 0 | 0 | 0 | 0 | 0 |
| 2020/11/2 | 0 | 0 | 0 | 0 | 0 | 0 | 0 | 0 | 0 | 0 |
| 2020/11/3 | 0 | 0 | 0 | 0 | 0 | 0 | 0 | 0 | 0 | 0 |
| 2020/11/4 | 1 | 0 | 0 | 0 | 0 | 0 | 0 | 0 | 0 | 100 |
| 2020/11/5 | 0 | 0 | 0 | 0 | 0 | 0 | 0 | 0 | 0 | 0 |
| 2020/11/6 | 2 | 0 | 0 | 0 | 0 | 0 | 0 | 0 | 0 | 100 |
| 2020/11/7 | 0 | 0 | 0 | 0 | 0 | 0 | 0 | 0 | 0 | 0 |
| 2020/11/8 | 0 | 0 | 0 | 0 | 0 | 0 | 0 | 0 | 0 | 0 |
| 2020/11/9 | 1 | 0 | 0 | 0 | 0 | 0 | 0 | 0 | 0 | 100 |
| 2020/11/10 | 1 | 0 | 0 | 0 | 0 | 0 | 0 | 0 | 0 | 100 |
| 2020/11/11 | 2 | 0 | 0 | 0 | 0 | 0 | 0 | 0 | 0 | 100 |
| 2020/11/12 | 0 | 0 | 0 | 0 | 0 | 0 | 0 | 0 | 0 | 0 |
| 2020/11/13 | 1 | 0 | 0 | 0 | 0 | 0 | 0 | 0 | 0 | 100 |
| 2020/11/14 | 1 | 0 | 0 | 0 | 0 | 0 | 0 | 0 | 0 | 100 |
| 2020/11/15 | 1 | 0 | 0 | 0 | 0 | 0 | 0 | 0 | 0 | 100 |
| 2020/11/16 | 1 | 0 | 0 | 0 | 0 | 0 | 0 | 0 | 0 | 100 |
| 2020/11/17 | 0 | 0 | 0 | 0 | 0 | 0 | 0 | 0 | 0 | 0 |
| 2020/11/18 | 0 | 0 | 0 | 0 | 0 | 0 | 0 | 0 | 0 | 0 |
| 2020/11/19 | 0 | 0 | 0 | 0 | 0 | 0 | 0 | 0 | 0 | 0 |
| 2020/11/20 | 1 | 0 | 0 | 0 | 0 | 0 | 0 | 0 | 0 | 100 |
| 2020/11/21 | 0 | 0 | 0 | 0 | 0 | 0 | 0 | 0 | 0 | 0 |
| 2020/11/22 | 1 | 0 | 0 | 0 | 0 | 0 | 0 | 0 | 0 | 100 |
| 2020/11/23 | 0 | 0 | 0 | 0 | 0 | 0 | 0 | 0 | 0 | 0 |
| 2020/11/24 | 0 | 0 | 0 | 0 | 0 | 0 | 0 | 0 | 0 | 0 |
| 2020/11/25 | 0 | 0 | 0 | 0 | 0 | 0 | 0 | 0 | 0 | 0 |
| 2020/11/26 | 5 | 0 | 0 | 0 | 0 | 0 | 0 | 0 | 0 | 100 |
| 2020/11/27 | 1 | 0 | 0 | 0 | 0 | 0 | 0 | 0 | 0 | 100 |
| 2020/11/28 | 0 | 0 | 0 | 0 | 0 | 0 | 0 | 0 | 0 | 0 |
| 2020/11/29 | 0 | 0 | 0 | 0 | 0 | 0 | 0 | 0 | 0 | 0 |
| 2020/11/30 | 1 | 0 | 0 | 0 | 0 | 0 | 0 | 0 | 0 | 100 |
| 2020/12/1 | 1 | 0 | 0 | 0 | 0 | 0 | 0 | 0 | 0 | 100 |
| 2020/12/2 | 2 | 0 | 0 | 0 | 0 | 0 | 0 | 0 | 0 | 100 |
| 2020/12/3 | 3 | 0 | 0 | 0 | 0 | 0 | 0 | 0 | 0 | 100 |
| 2020/12/4 | 2 | 0 | 0 | 0 | 0 | 0 | 0 | 0 | 0 | 100 |
| 2020/12/5 | 2 | 0 | 0 | 0 | 0 | 0 | 0 | 0 | 0 | 100 |
| 2020/12/6 | 1 | 0 | 0 | 0 | 0 | 0 | 0 | 0 | 0 | 100 |
| 2020/12/7 | 0 | 0 | 0 | 0 | 0 | 0 | 0 | 0 | 0 | 0 |
| 2020/12/8 | 3 | 0 | 0 | 0 | 0 | 0 | 0 | 0 | 0 | 100 |
| 2020/12/9 | 2 | 0 | 0 | 0 | 0 | 0 | 0 | 0 | 0 | 100 |
| 2020/12/10 | 0 | 0 | 0 | 0 | 0 | 0 | 0 | 0 | 0 | 0 |
| 2020/12/11 | 0 | 0 | 0 | 0 | 0 | 0 | 0 | 0 | 0 | 0 |

| | | | | | | | | | | |
|---|---|---|---|---|---|---|---|---|---|---|
| 2020/12/12 | 0 | 0 | 0 | 0 | 0 | 0 | 0 | 0 | 0 | 0 |
| 2020/12/13 | 0 | 0 | 0 | 0 | 0 | 0 | 0 | 0 | 0 | 0 |
| 2020/12/14 | 2 | 0 | 0 | 0 | 0 | 0 | 0 | 0 | 0 | 100 |
| 2020/12/15 | 3 | 0 | 0 | 0 | 0 | 0 | 0 | 0 | 0 | 100 |
| 2020/12/16 | 3 | 0 | 0 | 0 | 0 | 0 | 0 | 0 | 0 | 100 |
| 2020/12/17 | 5 | 0 | 0 | 0 | 0 | 0 | 0 | 0 | 0 | 100 |
| 2020/12/18 | 5 | 0 | 0 | 0 | 0 | 0 | 0 | 0 | 0 | 100 |
| 2020/12/19 | 2 | 0 | 0 | 0 | 0 | 0 | 0 | 0 | 0 | 100 |
| 2020/12/20 | 4 | 0 | 0 | 0 | 0 | 0 | 0 | 0 | 0 | 100 |
| 2020/12/21 | 6 | 0 | 0 | 0 | 0 | 0 | 0 | 0 | 0 | 100 |
| 2020/12/22 | 3 | 0 | 0 | 0 | 0 | 0 | 0 | 0 | 0 | 100 |
| 2020/12/23 | 3 | 0 | 0 | 0 | 0 | 0 | 0 | 0 | 0 | 100 |
| 2020/12/24 | 3 | 0 | 0 | 0 | 0 | 0 | 0 | 0 | 0 | 100 |
| 2020/12/25 | 1 | 0 | 0 | 0 | 0 | 0 | 0 | 0 | 0 | 100 |
| 2020/12/26 | 5 | 0 | 0 | 0 | 0 | 0 | 0 | 0 | 0 | 100 |
| 2020/12/27 | 9 | 0 | 0 | 0 | 0 | 0 | 0 | 0 | 0 | 100 |
| 2020/12/28 | 11 | 0 | 0 | 0 | 0 | 0 | 0 | 0 | 0 | 100 |
| 2020/12/29 | 4 | 0 | 0 | 0 | 0 | 0 | 0 | 0 | 0 | 100 |
| 2020/12/30 | 22 | 0 | 0 | 0 | 0 | 0 | 0 | 0 | 0 | 100 |
| 2020/12/31 | 2 | 0 | 0 | 0 | 0 | 0 | 0 | 0 | 0 | 100 |
| 2021/1/1 | 8 | 0 | 0 | 0 | 0 | 0 | 0 | 0 | 0 | 100 |
| 2021/1/2 | 6 | 0 | 0 | 0 | 0 | 0 | 0 | 0 | 0 | 100 |
| 2021/1/3 | 4 | 0 | 0 | 0 | 0 | 0 | 0 | 0 | 0 | 100 |
| 2021/1/4 | 7 | 1 | 0 | 0 | 0 | 14 | 0 | 0 | 0 | 86 |
| 2021/1/5 | 8 | 0 | 0 | 0 | 0 | 0 | 0 | 0 | 0 | 100 |
| 2021/1/6 | 6 | 1 | 0 | 0 | 0 | 17 | 0 | 0 | 0 | 83 |
| 2021/1/7 | 4 | 1 | 0 | 0 | 0 | 25 | 0 | 0 | 0 | 75 |
| 2021/1/8 | 8 | 0 | 0 | 0 | 0 | 0 | 0 | 0 | 0 | 100 |
| 2021/1/9 | 2 | 0 | 0 | 0 | 0 | 0 | 0 | 0 | 0 | 100 |
| 2021/1/10 | 10 | 1 | 0 | 0 | 0 | 10 | 0 | 0 | 0 | 90 |
| 2021/1/11 | 3 | 0 | 0 | 0 | 1 | 0 | 0 | 0 | 33 | 67 |
| 2021/1/12 | 9 | 1 | 0 | 0 | 0 | 11 | 0 | 0 | 0 | 89 |
| 2021/1/13 | 7 | 0 | 0 | 0 | 0 | 0 | 0 | 0 | 0 | 100 |
| 2021/1/14 | 8 | 0 | 0 | 0 | 0 | 0 | 0 | 0 | 0 | 100 |
| 2021/1/15 | 5 | 0 | 0 | 0 | 0 | 0 | 0 | 0 | 0 | 100 |
| 2021/1/16 | 7 | 0 | 0 | 0 | 0 | 0 | 0 | 0 | 0 | 100 |
| 2021/1/17 | 7 | 0 | 0 | 0 | 0 | 0 | 0 | 0 | 0 | 100 |
| 2021/1/18 | 10 | 0 | 0 | 0 | 0 | 0 | 0 | 0 | 0 | 100 |
| 2021/1/19 | 8 | 4 | 0 | 0 | 0 | 50 | 0 | 0 | 0 | 50 |
| 2021/1/20 | 9 | 1 | 0 | 0 | 0 | 11 | 0 | 0 | 0 | 89 |
| 2021/1/21 | 3 | 1 | 0 | 0 | 0 | 33 | 0 | 0 | 0 | 67 |
| 2021/1/22 | 5 | 2 | 0 | 0 | 0 | 40 | 0 | 0 | 0 | 60 |
| 2021/1/23 | 1 | 0 | 0 | 0 | 0 | 0 | 0 | 0 | 0 | 100 |
| 2021/1/24 | 1 | 0 | 0 | 0 | 0 | 0 | 0 | 0 | 0 | 100 |
| 2021/1/25 | 2 | 0 | 0 | 0 | 0 | 0 | 0 | 0 | 0 | 100 |
| 2021/1/26 | 3 | 0 | 0 | 0 | 0 | 0 | 0 | 0 | 0 | 100 |
| 2021/1/27 | 1 | 0 | 0 | 0 | 0 | 0 | 0 | 0 | 0 | 100 |
| 2021/1/28 | 1 | 0 | 0 | 0 | 0 | 0 | 0 | 0 | 0 | 100 |
| 2021/1/29 | 3 | 0 | 0 | 0 | 0 | 0 | 0 | 0 | 0 | 100 |
| 2021/1/30 | 7 | 2 | 0 | 0 | 0 | 29 | 0 | 0 | 0 | 71 |
| 2021/1/31 | 1 | 0 | 0 | 0 | 0 | 0 | 0 | 0 | 0 | 100 |
| 2021/2/1 | 1 | 1 | 0 | 0 | 0 | 100 | 0 | 0 | 0 | 0 |
| 2021/2/2 | 1 | 1 | 0 | 0 | 0 | 100 | 0 | 0 | 0 | 0 |
| 2021/2/3 | 1 | 0 | 0 | 0 | 0 | 0 | 0 | 0 | 0 | 100 |
| 2021/2/4 | 0 | 0 | 0 | 0 | 0 | 0 | 0 | 0 | 0 | 0 |
| 2021/2/5 | 5 | 1 | 0 | 0 | 0 | 20 | 0 | 0 | 0 | 80 |
| 2021/2/6 | 1 | 1 | 0 | 0 | 0 | 100 | 0 | 0 | 0 | 0 |

| | | | | | | | | | | |
|---|---|---|---|---|---|---|---|---|---|---|
| 2021/2/7 | 2 | 1 | 0 | 0 | 0 | 50 | 0 | 0 | 0 | 50 |
| 2021/2/8 | 0 | 0 | 0 | 0 | 0 | 0 | 0 | 0 | 0 | 0 |
| 2021/2/9 | 0 | 0 | 0 | 0 | 0 | 0 | 0 | 0 | 0 | 0 |
| 2021/2/10 | 1 | 0 | 0 | 0 | 0 | 0 | 0 | 0 | 0 | 100 |
| 2021/2/11 | 1 | 0 | 0 | 0 | 0 | 0 | 0 | 0 | 0 | 100 |
| 2021/2/12 | 4 | 1 | 0 | 0 | 0 | 25 | 0 | 0 | 0 | 75 |
| 2021/2/13 | 2 | 0 | 0 | 0 | 0 | 0 | 0 | 0 | 0 | 100 |
| 2021/2/14 | 2 | 1 | 0 | 0 | 0 | 50 | 0 | 0 | 0 | 50 |
| 2021/2/15 | 2 | 0 | 0 | 0 | 0 | 0 | 0 | 0 | 0 | 100 |
| 2021/2/16 | 0 | 0 | 0 | 0 | 0 | 0 | 0 | 0 | 0 | 0 |
| 2021/2/17 | 2 | 1 | 0 | 0 | 0 | 50 | 0 | 0 | 0 | 50 |
| 2021/2/18 | 2 | 2 | 0 | 0 | 0 | 100 | 0 | 0 | 0 | 0 |
| 2021/2/19 | 4 | 1 | 0 | 0 | 0 | 25 | 0 | 0 | 0 | 75 |
| 2021/2/20 | 1 | 1 | 0 | 0 | 0 | 100 | 0 | 0 | 0 | 0 |
| 2021/2/21 | 1 | 0 | 0 | 0 | 1 | 0 | 0 | 0 | 100 | 0 |
| 2021/2/22 | 1 | 0 | 0 | 0 | 0 | 0 | 0 | 0 | 0 | 100 |
| 2021/2/23 | 2 | 0 | 0 | 0 | 0 | 0 | 0 | 0 | 0 | 100 |
| 2021/2/24 | 1 | 0 | 0 | 0 | 0 | 0 | 0 | 0 | 0 | 100 |
| 2021/2/25 | 4 | 0 | 0 | 0 | 0 | 0 | 0 | 0 | 0 | 100 |
| 2021/2/26 | 7 | 0 | 0 | 0 | 3 | 0 | 0 | 0 | 43 | 57 |
| 2021/2/27 | 3 | 0 | 0 | 0 | 2 | 0 | 0 | 0 | 67 | 33 |
| 2021/2/28 | 3 | 0 | 0 | 0 | 0 | 0 | 0 | 0 | 0 | 100 |
| 2021/3/1 | 8 | 0 | 0 | 0 | 0 | 0 | 0 | 0 | 0 | 100 |
| 2021/3/2 | 4 | 0 | 0 | 0 | 1 | 0 | 0 | 0 | 25 | 75 |
| 2021/3/3 | 7 | 1 | 0 | 0 | 3 | 14 | 0 | 0 | 43 | 43 |
| 2021/3/4 | 6 | 2 | 0 | 0 | 2 | 33 | 0 | 0 | 33 | 33 |
| 2021/3/5 | 11 | 0 | 0 | 0 | 1 | 0 | 0 | 0 | 9 | 91 |
| 2021/3/6 | 8 | 1 | 0 | 0 | 1 | 13 | 0 | 0 | 13 | 75 |
| 2021/3/7 | 4 | 0 | 0 | 0 | 0 | 0 | 0 | 0 | 0 | 100 |
| 2021/3/8 | 14 | 1 | 0 | 0 | 4 | 7 | 0 | 0 | 29 | 64 |
| 2021/3/9 | 3 | 1 | 0 | 0 | 1 | 33 | 0 | 0 | 33 | 33 |
| 2021/3/10 | 10 | 3 | 0 | 0 | 5 | 30 | 0 | 0 | 50 | 20 |
| 2021/3/11 | 2 | 0 | 0 | 0 | 1 | 0 | 0 | 0 | 50 | 50 |
| 2021/3/12 | 7 | 2 | 0 | 0 | 1 | 29 | 0 | 0 | 14 | 57 |
| 2021/3/13 | 3 | 1 | 0 | 0 | 0 | 33 | 0 | 0 | 0 | 67 |
| 2021/3/14 | 4 | 0 | 0 | 0 | 1 | 0 | 0 | 0 | 25 | 75 |
| 2021/3/15 | 7 | 1 | 0 | 0 | 3 | 14 | 0 | 0 | 43 | 43 |
| 2021/3/16 | 3 | 0 | 0 | 0 | 0 | 0 | 0 | 0 | 0 | 100 |
| 2021/3/17 | 6 | 2 | 0 | 0 | 1 | 33 | 0 | 0 | 17 | 50 |
| 2021/3/18 | 8 | 1 | 0 | 0 | 1 | 13 | 0 | 0 | 13 | 75 |
| 2021/3/19 | 4 | 2 | 0 | 0 | 1 | 50 | 0 | 0 | 25 | 25 |
| 2021/3/20 | 3 | 2 | 0 | 0 | 0 | 67 | 0 | 0 | 0 | 33 |
| 2021/3/21 | 3 | 0 | 0 | 0 | 1 | 0 | 0 | 0 | 33 | 67 |
| 2021/3/22 | 3 | 0 | 0 | 0 | 0 | 0 | 0 | 0 | 0 | 100 |
| 2021/3/23 | 6 | 1 | 0 | 0 | 3 | 17 | 0 | 0 | 50 | 33 |
| 2021/3/24 | 13 | 1 | 0 | 0 | 3 | 8 | 0 | 0 | 23 | 69 |
| 2021/3/25 | 6 | 1 | 0 | 0 | 1 | 17 | 0 | 0 | 17 | 67 |
| 2021/3/26 | 8 | 1 | 0 | 0 | 2 | 13 | 0 | 0 | 25 | 63 |
| 2021/3/27 | 8 | 1 | 0 | 0 | 1 | 13 | 0 | 0 | 13 | 75 |
| 2021/3/28 | 2 | 0 | 0 | 0 | 1 | 0 | 0 | 0 | 50 | 50 |
| 2021/3/29 | 7 | 2 | 0 | 0 | 0 | 29 | 0 | 0 | 0 | 71 |
| 2021/3/30 | 5 | 0 | 0 | 1 | 0 | 0 | 0 | 20 | 0 | 80 |
| 2021/3/31 | 9 | 2 | 0 | 0 | 2 | 22 | 0 | 0 | 22 | 56 |
| 2021/4/1 | 12 | 2 | 0 | 0 | 5 | 17 | 0 | 0 | 42 | 42 |
| 2021/4/2 | 8 | 0 | 0 | 0 | 3 | 0 | 0 | 0 | 38 | 63 |
| 2021/4/3 | 18 | 9 | 0 | 0 | 6 | 50 | 0 | 0 | 33 | 17 |
| 2021/4/4 | 12 | 1 | 0 | 0 | 7 | 8 | 0 | 0 | 58 | 33 |

| | | | | | | | | | | |
|---|---|---|---|---|---|---|---|---|---|---|
| 2021/4/5 | 84 | 32 | 0 | 2 | 17 | 38 | 0 | 2 | 20 | 39 |
| 2021/4/6 | 36 | 11 | 0 | 0 | 11 | 31 | 0 | 0 | 31 | 39 |
| 2021/4/7 | 37 | 18 | 0 | 0 | 8 | 49 | 0 | 0 | 22 | 30 |
| 2021/4/8 | 71 | 18 | 0 | 0 | 20 | 25 | 0 | 0 | 28 | 46 |
| 2021/4/9 | 73 | 20 | 0 | 6 | 18 | 27 | 0 | 8 | 25 | 40 |
| 2021/4/10 | 42 | 8 | 0 | 0 | 7 | 19 | 0 | 0 | 17 | 64 |
| 2021/4/11 | 13 | 4 | 0 | 0 | 4 | 31 | 0 | 0 | 31 | 38 |
| 2021/4/12 | 55 | 14 | 0 | 3 | 11 | 25 | 0 | 5 | 20 | 49 |
| 2021/4/13 | 43 | 12 | 0 | 0 | 14 | 28 | 0 | 0 | 33 | 40 |
| 2021/4/14 | 46 | 24 | 0 | 1 | 8 | 52 | 0 | 2 | 17 | 28 |
| 2021/4/15 | 55 | 26 | 0 | 1 | 16 | 47 | 0 | 2 | 29 | 22 |
| 2021/4/16 | 29 | 11 | 0 | 1 | 8 | 38 | 0 | 3 | 28 | 31 |
| 2021/4/17 | 13 | 6 | 0 | 0 | 3 | 46 | 0 | 0 | 23 | 31 |
| 2021/4/18 | 7 | 2 | 0 | 0 | 2 | 29 | 0 | 0 | 29 | 43 |
| 2021/4/19 | 23 | 8 | 0 | 0 | 5 | 35 | 0 | 0 | 22 | 43 |
| 2021/4/20 | 18 | 5 | 0 | 1 | 8 | 28 | 0 | 6 | 44 | 22 |
| 2021/4/21 | 30 | 6 | 0 | 1 | 13 | 20 | 0 | 3 | 43 | 33 |
| 2021/4/22 | 20 | 9 | 0 | 0 | 9 | 45 | 0 | 0 | 45 | 10 |
| 2021/4/23 | 13 | 2 | 0 | 0 | 8 | 15 | 0 | 0 | 62 | 23 |
| 2021/4/24 | 13 | 4 | 0 | 0 | 4 | 31 | 0 | 0 | 31 | 38 |
| 2021/4/25 | 11 | 2 | 0 | 1 | 7 | 18 | 0 | 9 | 64 | 9 |
| 2021/4/26 | 39 | 10 | 0 | 1 | 16 | 26 | 0 | 3 | 41 | 31 |
| 2021/4/27 | 32 | 10 | 0 | 0 | 6 | 31 | 0 | 0 | 19 | 50 |
| 2021/4/28 | 18 | 5 | 0 | 4 | 3 | 28 | 0 | 22 | 17 | 33 |
| 2021/4/29 | 20 | 12 | 0 | 0 | 3 | 60 | 0 | 0 | 15 | 25 |
| 2021/4/30 | 21 | 7 | 0 | 0 | 7 | 33 | 0 | 0 | 33 | 33 |
| 2021/5/1 | 15 | 4 | 0 | 0 | 7 | 27 | 0 | 0 | 47 | 27 |
| 2021/5/2 | 7 | 2 | 0 | 0 | 5 | 29 | 0 | 0 | 71 | 0 |
| 2021/5/3 | 34 | 13 | 0 | 1 | 11 | 38 | 0 | 3 | 32 | 26 |
| 2021/5/4 | 21 | 4 | 0 | 2 | 10 | 19 | 0 | 10 | 48 | 24 |
| 2021/5/5 | 23 | 6 | 0 | 0 | 12 | 26 | 0 | 0 | 52 | 22 |
| 2021/5/6 | 21 | 5 | 0 | 1 | 11 | 24 | 0 | 5 | 52 | 19 |
| 2021/5/7 | 28 | 6 | 0 | 0 | 16 | 21 | 0 | 0 | 57 | 21 |
| 2021/5/8 | 31 | 5 | 0 | 1 | 14 | 16 | 0 | 3 | 45 | 35 |
| 2021/5/9 | 7 | 1 | 0 | 0 | 4 | 14 | 0 | 0 | 57 | 29 |
| 2021/5/10 | 29 | 10 | 0 | 0 | 10 | 34 | 0 | 0 | 34 | 31 |
| 2021/5/11 | 13 | 3 | 0 | 2 | 7 | 23 | 0 | 15 | 54 | 8 |
| 2021/5/12 | 23 | 4 | 0 | 0 | 14 | 17 | 0 | 0 | 61 | 22 |
| 2021/5/13 | 31 | 11 | 0 | 1 | 9 | 35 | 0 | 3 | 29 | 32 |
| 2021/5/14 | 14 | 3 | 0 | 1 | 8 | 21 | 0 | 7 | 57 | 14 |
| 2021/5/15 | 10 | 5 | 0 | 1 | 4 | 50 | 0 | 10 | 40 | 0 |
| 2021/5/16 | 7 | 1 | 0 | 0 | 4 | 14 | 0 | 0 | 57 | 29 |
| 2021/5/17 | 5 | 3 | 0 | 0 | 2 | 60 | 0 | 0 | 40 | 0 |
| 2021/5/18 | 25 | 4 | 0 | 0 | 14 | 16 | 0 | 0 | 56 | 28 |
| 2021/5/19 | 32 | 12 | 0 | 0 | 17 | 38 | 0 | 0 | 53 | 9 |
| 2021/5/20 | 17 | 6 | 0 | 0 | 11 | 35 | 0 | 0 | 65 | 0 |
| 2021/5/21 | 12 | 5 | 0 | 0 | 5 | 42 | 0 | 0 | 42 | 17 |
| 2021/5/22 | 1 | 0 | 0 | 0 | 1 | 0 | 0 | 0 | 100 | 0 |
| 2021/5/23 | 3 | 2 | 0 | 0 | 1 | 67 | 0 | 0 | 33 | 0 |
| 2021/5/24 | 9 | 3 | 0 | 0 | 5 | 33 | 0 | 0 | 56 | 11 |
| 2021/5/25 | 6 | 3 | 0 | 0 | 1 | 50 | 0 | 0 | 17 | 33 |
| 2021/5/26 | 5 | 1 | 0 | 1 | 3 | 20 | 0 | 20 | 60 | 0 |
| 2021/5/27 | 3 | 0 | 0 | 0 | 3 | 0 | 0 | 0 | 100 | 0 |
| 2021/5/28 | 2 | 1 | 0 | 0 | 1 | 50 | 0 | 0 | 50 | 0 |
| 2021/5/29 | 5 | 2 | 0 | 0 | 3 | 40 | 0 | 0 | 60 | 0 |
| 2021/5/30 | 2 | 0 | 0 | 0 | 2 | 0 | 0 | 0 | 100 | 0 |
| 2021/5/31 | 0 | 0 | 0 | 0 | 0 | 0 | 0 | 0 | 0 | 0 |

| | | | | | | | | | | |
|---|---|---|---|---|---|---|---|---|---|---|
| 2021/6/1 | 3 | 1 | 0 | 0 | 2 | 33 | 0 | 0 | 67 | 0 |
| 2021/6/2 | 3 | 0 | 0 | 0 | 2 | 0 | 0 | 0 | 67 | 33 |
| 2021/6/3 | 4 | 1 | 0 | 0 | 3 | 25 | 0 | 0 | 75 | 0 |
| 2021/6/4 | 11 | 0 | 0 | 0 | 6 | 0 | 0 | 0 | 55 | 45 |
| 2021/6/5 | 2 | 1 | 0 | 0 | 1 | 50 | 0 | 0 | 50 | 0 |
| 2021/6/6 | 1 | 0 | 0 | 0 | 1 | 0 | 0 | 0 | 100 | 0 |
| 2021/6/7 | 3 | 0 | 0 | 0 | 3 | 0 | 0 | 0 | 100 | 0 |
| 2021/6/8 | 13 | 3 | 0 | 1 | 7 | 23 | 0 | 8 | 54 | 15 |
| 2021/6/9 | 6 | 1 | 0 | 0 | 5 | 17 | 0 | 0 | 83 | 0 |
| 2021/6/10 | 10 | 3 | 0 | 1 | 5 | 30 | 0 | 10 | 50 | 10 |
| 2021/6/11 | 10 | 0 | 0 | 0 | 8 | 0 | 0 | 0 | 80 | 20 |
| 2021/6/12 | 6 | 2 | 0 | 0 | 2 | 33 | 0 | 0 | 33 | 33 |
| 2021/6/13 | 5 | 1 | 0 | 1 | 3 | 20 | 0 | 20 | 60 | 0 |
| 2021/6/14 | 6 | 2 | 0 | 0 | 3 | 33 | 0 | 0 | 50 | 17 |
| 2021/6/15 | 12 | 3 | 0 | 0 | 9 | 25 | 0 | 0 | 75 | 0 |
| 2021/6/16 | 5 | 0 | 0 | 0 | 5 | 0 | 0 | 0 | 100 | 0 |
| 2021/6/17 | 4 | 1 | 0 | 0 | 2 | 25 | 0 | 0 | 50 | 25 |
| 2021/6/18 | 1 | 0 | 0 | 0 | 1 | 0 | 0 | 0 | 100 | 0 |
| 2021/6/19 | 0 | 0 | 0 | 0 | 0 | 0 | 0 | 0 | 0 | 0 |
| 2021/6/20 | 2 | 0 | 0 | 0 | 2 | 0 | 0 | 0 | 100 | 0 |
| 2021/6/21 | 10 | 0 | 0 | 1 | 8 | 0 | 0 | 10 | 80 | 10 |
| 2021/6/22 | 4 | 0 | 0 | 0 | 4 | 0 | 0 | 0 | 100 | 0 |
| 2021/6/23 | 7 | 5 | 0 | 0 | 2 | 71 | 0 | 0 | 29 | 0 |
| 2021/6/24 | 7 | 2 | 0 | 0 | 5 | 29 | 0 | 0 | 71 | 0 |
| 2021/6/25 | 0 | 0 | 0 | 0 | 0 | 0 | 0 | 0 | 0 | 0 |
| 2021/6/26 | 1 | 0 | 0 | 0 | 1 | 0 | 0 | 0 | 100 | 0 |
| 2021/6/27 | 0 | 0 | 0 | 0 | 0 | 0 | 0 | 0 | 0 | 0 |
| 2021/6/28 | 10 | 0 | 0 | 0 | 9 | 0 | 0 | 0 | 90 | 10 |
| 2021/6/29 | 10 | 0 | 0 | 2 | 8 | 0 | 0 | 20 | 80 | 0 |
| 2021/6/30 | 15 | 3 | 0 | 0 | 11 | 20 | 0 | 0 | 73 | 7 |
| 2021/7/1 | 11 | 0 | 0 | 1 | 8 | 0 | 0 | 9 | 73 | 18 |
| 2021/7/2 | 6 | 1 | 0 | 0 | 4 | 17 | 0 | 0 | 67 | 17 |
| 2021/7/3 | 8 | 0 | 1 | 1 | 6 | 0 | 13 | 13 | 75 | 0 |
| 2021/7/4 | 3 | 0 | 0 | 0 | 3 | 0 | 0 | 0 | 100 | 0 |
| 2021/7/5 | 4 | 0 | 0 | 0 | 3 | 0 | 0 | 0 | 75 | 25 |
| 2021/7/6 | 6 | 0 | 0 | 0 | 5 | 0 | 0 | 0 | 83 | 17 |
| 2021/7/7 | 11 | 1 | 1 | 0 | 9 | 9 | 9 | 0 | 82 | 0 |
| 2021/7/8 | 7 | 1 | 1 | 0 | 3 | 14 | 14 | 0 | 43 | 29 |
| 2021/7/9 | 7 | 0 | 0 | 0 | 6 | 0 | 0 | 0 | 86 | 14 |
| 2021/7/10 | 4 | 0 | 0 | 1 | 3 | 0 | 0 | 25 | 75 | 0 |
| 2021/7/11 | 1 | 0 | 0 | 0 | 1 | 0 | 0 | 0 | 100 | 0 |
| 2021/7/12 | 4 | 0 | 0 | 0 | 3 | 0 | 0 | 0 | 75 | 25 |
| 2021/7/13 | 7 | 0 | 0 | 0 | 7 | 0 | 0 | 0 | 100 | 0 |
| 2021/7/14 | 15 | 1 | 0 | 1 | 11 | 7 | 0 | 7 | 73 | 13 |
| 2021/7/15 | 11 | 1 | 0 | 0 | 9 | 9 | 0 | 0 | 82 | 9 |
| 2021/7/16 | 9 | 0 | 5 | 0 | 3 | 0 | 56 | 0 | 33 | 11 |
| 2021/7/17 | 4 | 0 | 0 | 0 | 4 | 0 | 0 | 0 | 100 | 0 |
| 2021/7/18 | 2 | 0 | 0 | 0 | 2 | 0 | 0 | 0 | 100 | 0 |
| 2021/7/19 | 6 | 0 | 1 | 0 | 5 | 0 | 17 | 0 | 83 | 0 |
| 2021/7/20 | 7 | 0 | 0 | 1 | 6 | 0 | 0 | 14 | 86 | 0 |
| 2021/7/21 | 5 | 0 | 0 | 0 | 5 | 0 | 0 | 0 | 100 | 0 |
| 2021/7/22 | 2 | 0 | 0 | 0 | 2 | 0 | 0 | 0 | 100 | 0 |
| 2021/7/23 | 5 | 0 | 0 | 0 | 5 | 0 | 0 | 0 | 100 | 0 |
| 2021/7/24 | 5 | 0 | 2 | 0 | 3 | 0 | 40 | 0 | 60 | 0 |
| 2021/7/25 | 0 | 0 | 0 | 0 | 0 | 0 | 0 | 0 | 0 | 0 |
| 2021/7/26 | 2 | 0 | 1 | 0 | 1 | 0 | 50 | 0 | 50 | 0 |
| 2021/7/27 | 1 | 0 | 0 | 0 | 1 | 0 | 0 | 0 | 100 | 0 |

| | | | | | | | | | | |
|---|---|---|---|---|---|---|---|---|---|---|
| 2021/7/28 | 2 | 0 | 1 | 0 | 1 | 0 | 50 | 0 | 50 | 0 |
| 2021/7/29 | 1 | 0 | 1 | 0 | 0 | 0 | 100 | 0 | 0 | 0 |
| 2021/7/30 | 0 | 0 | 0 | 0 | 0 | 0 | 0 | 0 | 0 | 0 |
| 2021/7/31 | 0 | 0 | 0 | 0 | 0 | 0 | 0 | 0 | 0 | 0 |
| 2021/8/1 | 0 | 0 | 0 | 0 | 0 | 0 | 0 | 0 | 0 | 0 |
| 2021/8/2 | 0 | 0 | 0 | 0 | 0 | 0 | 0 | 0 | 0 | 0 |
| 2021/8/3 | 0 | 0 | 0 | 0 | 0 | 0 | 0 | 0 | 0 | 0 |
| 2021/8/4 | 1 | 0 | 0 | 0 | 1 | 0 | 0 | 0 | 100 | 0 |

**Table S3. Haplotypes of the spike protein of Mu variant.**

| Haplotype | Sequence number | Percentage |
|---|---|---|
| T95I\|YY144-145TSN\|R346K\|E484K\|N501Y\|D614G\|P681H\|D950N | 1488 | 39.6 |
| T95I\|YY144-145TSN\|R346K\|K417N\|E484K\|N501Y\|D614G\|P681H\|D950N | 189 | 5.0 |
| T95I\|YY144-145TSN\|R346K\|E484K\|N501Y\|D614G\|P681H | 129 | 3.4 |
| T95I\|YY144-145TSN\|R346K\|E484K\|N501Y\|D614G\|P681H\|D950N\|M1229I | 128 | 3.4 |
| T95I\|YY144-145SN\|R346K\|E484K\|N501Y\|D614G\|P681H\|D950N | 111 | 3.0 |
| T95I\|V143VT\|Y145N\|R346K\|E484K\|N501Y\|D614G\|P681H\|D950N | 108 | 2.9 |
| T95I\|R346K\|E484K\|N501Y\|D614G\|P681H\|D950N | 76 | 2.0 |
| T95I\|YY144-145TSN\|R346K\|E484K\|N501Y\|T572I\|D614G\|P681H\|D950N | 58 | 1.5 |
| L5F\|T95I\|YY144-145TSN\|R346K\|E484K\|N501Y\|D614G\|P681H\|D950N | 45 | 1.2 |
| T95I\|YY144-145TSN\|R346K\|D614G\|P681H\|D950N | 43 | 1.1 |

**Table S4. Mutations in the spike protein of Mu variant.**

| Mutation | Sequence number | Percentage |
|---|---|---|
| D614G | 3,755 | 99.8 |
| P681H | 3,743 | 99.5 |
| R346K | 3,684 | 97.9 |
| N501Y | 3,607 | 95.9 |
| T95I | 3,604 | 95.8 |
| E484K | 3,591 | 95.5 |
| D950N | 3,385 | 90.0 |
| Y145N | 3,264 | 86.8 |
| Y144S | 3,053 | 81.2 |
| V143VT | 2,952 | 78.5 |
| K417N | 302 | 8.0 |
| M1229I | 192 | 5.1 |
| Y144T | 97 | 2.6 |
| T572I | 96 | 2.6 |
| Y145SN | 71 | 1.9 |
| M1237I | 64 | 1.7 |
| L5F | 64 | 1.7 |
| E583D | 41 | 1.1 |
| E1258D | 39 | 1.0 |

**Table S5. Mutations in the spike proteins of SARS-CoV-2 variants used in this study.**

| Classification | Parental | Alpha | Beta | Gamma | Delta | Epsilon | Lambda | Mu |
|---|---|---|---|---|---|---|---|---|
| PANGO lineage | B.1 | B.1.1.7 | B.1.351 | P.1 | B.1.617.2 | B.1.427 | C.37 | B.1.621 |
| Mutations | D614G | HV69-70del | L18F | L18F | T19R | S13I | G75V | T95I |
| | | Y144del | D80A | T20N | G142D | W152C | T76I | YY144-145TSN |
| | | N501Y | D215G | P26S | EFR156-158G | L452R | RSYLTPGD246-253N | R346K |
| | | A570D | R246I | D138Y | L452R | D614G | L452Q | E484K |
| | | D614G | K417N | R190S | T478K | | F490S | N501Y |
| | | P681H | E484K | K417T | D614G | | D614G | D614G |
| | | T716I | N501Y | E484K | P681R | | T859N | P681H |
| | | S982A | D614G | N501Y | D950N | | | D950N |
| | | D1118H | A701V | D614G | | | | |
| | | | | H655Y | | | | |
| | | | | T1027I | | | | |

**Table S6. Summary of COVID-19 convalescent sera used in this study.**

| Donor ID | Sex | Age | Severity | Date of test (MM/DD/YY) | Date of sampling (MM/DD/YY) | $NT_{50}$ | | | | | | | |
|---|---|---|---|---|---|---|---|---|---|---|---|---|---|
| | | | | | | Parental | Alpha | Beta | Gamma | Delta | Epsilon | Lambda | Mu |
| 01-07 | Male | 57 | Severe | 09/01/20 | 11/04/20 | 1,609 | 890 | 139 | 256 | 482 | 640 | 414 | 94 |
| 03-03 | Male | 58 | Severe | 09/30/20 | 12/04/20 | 500 | 158 | <40 | 131 | 92 | 200 | 163 | <40 |
| 12-01 | Male | 86 | Moderate | 08/19/20 | 09/18/20 | 293 | 370 | 117 | 95 | 222 | 260 | 159 | 44 |
| PS315 [a] | Female | 61 | NA [b] | 04/01/20 | 05/03/20 | 718 | 193 | 63 | 166 | 98 | 145 | 247 | <40 |
| PS324 [a] | Female | 76 | NA [b] | 04/01/20 | 05/03/20 | 773 | 340 | 84 | 155 | 194 | 362 | 217 | 106 |
| PS329 [a] | Female | 92 | NA [b] | 04/02/20 | 05/03/20 | 7,834 | 2,801 | 1,365 | 2,697 | 5,039 | 3,613 | 4,442 | 844 |
| PS362 [a] | Female | 52 | NA [b] | 04/17/20 | 05/17/20 | 834 | 235 | 194 | 376 | 199 | 404 | 174 | 46 |
| PS364 [a] | Female | 74 | NA [b] | 04/17/20 | 05/17/20 | 2,579 | 893 | 320 | 537 | 503 | 624 | 489 | 260 |
| | | | | | Geometric mean | 1,104 | 460 | 184 | 286 | 314 | 447 | 348 | 128 |

[a] Purchased from RayBiotech.

[b] Not applicable.

**Table S7. Summary of BNT162b2-vaccinated sera used in this study.**

| | | | Date of 2nd vaccination | Date of sampling | 50% neutralization titer | | | | | | | |
|---|---|---|---|---|---|---|---|---|---|---|---|---|
| Donor ID | Sex | Age | (MM/DD/YY) | (MM/DD/YY) | Parental | Alpha | Beta | Gamma | Delta | Epsilon | Lambda | Mu |
| #005-3 | Female | 34 | 04/30/21 | 05/27/21 | 555 | 440 | 64 | 168 | 84 | 225 | 228 | <40 |
| #007-3 | Female | 35 | 04/26/21 | 05/24/21 | 613 | 415 | 128 | 202 | 260 | 138 | 219 | <40 |
| #021-3 | Female | 47 | 04/27/21 | 05/25/21 | 361 | 161 | 52 | 73 | 151 | 122 | 237 | 46 |
| #026-3 | Female | 28 | 04/26/21 | 05/25/21 | 206 | 179 | 58 | 289 | 149 | 581 | 127 | 69 |
| #028-3 | Female | 42 | 04/28/21 | 05/26/21 | 595 | 597 | <40 | 112 | 520 | 463 | 434 | 112 |
| #030-3 | Female | 29 | 04/27/21 | 05/25/21 | 256 | 81 | 97 | 112 | 224 | 319 | 272 | 59 |
| #045-3 | Female | 34 | 04/30/21 | 05/24/21 | 715 | 271 | 133 | 323 | 239 | 276 | 354 | 137 |
| #056-3 | Male | 38 | 04/30/21 | 05/26/21 | 854 | 420 | 286 | 535 | 349 | 274 | 578 | 255 |
| #078-3 | Female | 32 | 04/22/21 | 05/06/21 | 1,260 | 691 | 141 | 291 | 265 | 347 | 579 | 121 |
| #103-3 | Male | 35 | 04/07/21 | 05/07/21 | 313 | 1,407 | 66 | 112 | 511 | 203 | 406 | 44 |
| | | | | Geometric mean | 497 | 351 | 98 | 186 | 240 | 265 | 311 | 89 |

**Table S8. Primers used for the construction of Mu spike expression plasmid.**

| Primer name | Sequence (5'-to-3') |
|---|---|
| T95I Fwd | TCTACTTTGCCAGCAttGAGAAGAGCAACATC |
| T95I Rev | GATGTTGCTCTTCTCaaTGCTGGCAAAGTAGA |
| YY144-145TSN Fwd | CCATTCCTGGGAGTCacctccaacCACAAGAACAACAAG |
| YY144-145TSN Rev | CTTGTTGTTCTTGTGgttggaggtGACTCCCAGGAATGG |
| R346K Fwd | GTTCAATGCCACCAaGTTTGCCTCTGTCT |
| R346K Rev | AGACAGAGGCAAACtTGGTGGCATTGAAC |
| E484K Fwd | CCATGTAATGGAGTGaAGGGCTTCAACTGTT |
| E484K Rev | AACAGTTGAAGCCCTtCACTCCATTACATGG |
| N501Y Fwd | TGGCTTCCAACCAACCtATGGAGTGGGCTA |
| N501Y Rev | TAGCCCACTCCATaGGTTGGTTGGAAGCCA |
| P681H Fwd | CCCAGACCAACAGCCatAGGAGGGCAAGGTCT |
| P681H Rev | AGACCTTGCCCTCCTatGGCTGTTGGTCTGGG |
| D950N Fwd | CTGGGCAAACTCCAAaATGTGGTGAACCAG |
| D950N Rev | CTGGTTCACCACATtTTGGAGTTTGCCCAG |

### Consortia

#### The Genotype to Phenotype Japan (G2P-Japan) Consortium

**Institute of Medical Science, University of Tokyo, Japan**
Jumpei Ito, Daichi Yamasoba, Yusuke Kosugi, Mai Suganami, Akiko Oide, Miyabishara Yokoyama, Mika Chiba

**Tokai University, Japan**
Jiaqi Wu, Miyoko Takahashi

**Kyoto University, Japan**
Yasuhiro Kazuma, Ryosuke Nomura, Yoshihito Horisawa

**Chiba University, Japan**
Motoaki Seki, Ryoji Fujiki, Tadanaga Shimada, Kiyoshi Hirahara, Koutaro Yokote, Toshinori Nakayama

**Hiroshima University, Japan**
Takashi Irie, Ryoko Kawabata, Nanami Morizako

**Hokkaido University, Japan**
Takasuke Fukuhara, Kenta Shimizu, Kana Tsushima, Haruko Kubo

**Kumamoto University, Japan**
Terumasa Ikeda, Chihiro Motozono, Hesham Nasser, Ryo Shimizu, Yue Yuan, Kazuko Kitazato, Haruyo Hasebe, Takamasa Ueno

**University of Miyazaki, Japan**
Akatsuki Saito, Erika P Butlertanaka, Yuri L Tanaka

**National Institute of Infectious Diseases, Japan**
Kenzo Tokunaga, Seiya Ozono

**Tokyo Metropolitan Institute of Public Health, Japan**
Kenji Sadamasu, Hiroyuki Asakura, Isao Yoshida, Mami Nagashima, Kazuhisa Yoshimura

**Acknowledgments**

We would like to thank all members of The Genotype to Phenotype Japan (G2P-Japan) Consortium. The super-computing resource was provided by the Human Genome Center at the University of Tokyo and the NIG supercomputer at ROIS National Institute of Genetics. We thank Dr. Kenzo Tokunaga (National Institute of Infectious Diseases, Japan) for sharing materials, Dr. Daniel Sauter (University Hospital Tübingen, Germany) for proofreading and helpful comments, and Dr. Paúl Cárdenas (Universidad San Francisco de Quito, Ecuador) for helpful suggestions.

This study was supported in part by AMED Research Program on Emerging and Re-emerging Infectious Diseases 20fk0108146 (to Kei Sato), 20fk0108270 (to Kei Sato), 20fk0108413 (to Atsushi Kaneda, So Nakagawa and Kei Sato) and 20fk0108451 (to G2P-Japan Consortium, Akifumi Takaori-Kondo, Atsushi Kaneda, So Nakagawa and Kei Sato); AMED Research Program on HIV/AIDS 21fk0410039 (to Kotaro Shirakawa and Kei Sato); JST SICORP (e-ASIA) JPMJSC20U1 (to Kei Sato); JST SICORP JPMJSC21U5 (to Kei Sato), JST CREST JPMJCR20H6 (to So Nakagawa) and JPMJCR20H4 (to Atsushi Kaneda and Kei Sato); JSPS KAKENHI Grants-in-Aid for Scientific Research B 18H02662 (to Kei Sato) and 21H02737 (to Kei Sato); JSPS Fund for the Promotion of Joint International Research (Fostering Joint International Research) 18KK0447 (to Kei Sato); JSPS Core-to-Core Program JPJSCCA20190008 (A. Advanced Research Networks) (to Kei Sato); JSPS Research Fellow DC1 19J20488 (to Izumi Kimura); The Tokyo Biochemical Research Foundation (to Kei Sato); and Joint Usage/Research Center program of Institute for Frontier Life and Medical Sciences, Kyoto University (to Kei Sato).
